# CPS: Mapping Physical Coordinates to High-Fidelity Spatial Transcriptomics via Privileged Multi-Scale Context Distillation

**DOI:** 10.64898/2026.04.05.716532

**Authors:** Lei Zhang, Kai Cao, Shuqiao Zheng, Shu Liang, Lin Wan

## Abstract

**Motivation:** Spatial transcriptomics enables the dissection of tissue heterogeneity within native contexts, yet current platforms are inherently constrained by high sparsity and low signal-to-noise ratios that obscure fine-grained biological signals. Current efforts to recover these signals are limited by image registration dependencies or the inherent context-blindness of implicit neural representations.

**Results:** We introduce the Cell Positioning System (CPS), a context-aware implicit neural representation framework designed to map physical coordinates to high-fidelity spatial transcriptomics via a privileged multi-scale context distillation strategy. CPS treats multi-scale tissue niches as privileged information, employing a teacher network equipped with a multi-scale niche attention mechanism to capture adaptive biological interactions during training. This structural knowledge is explicitly distilled into a student coordinate network, enabling the generation of context-aware expression landscapes solely from spatial coordinates during inference. Benchmarking on the DLPFC dataset demonstrates that CPS achieves state-of-the-art performance in spatial and gene expression imputation and denoising. Furthermore, CPS enables super-resolution to recover high-resolution mouse brain anatomical details and offers interpretability by identifying the scale effective size of biological interactions within human breast cancer tissues. Finally, the framework exhibits superior scalability for large-scale datasets with linear computational complexity.

**Availability:** Software is available online at https://github.com/tju-zl/CPS.

## 1. Introduction

Spatial transcriptomics (ST) has emerged as a transformative technology for dissecting tissue heterogeneity by mapping gene expression within its native tissue context[1]. However, the data quality of current platforms, such as 10x Visium and Stereo-seq, is inherently constrained by technical limitations, particularly high sparsity and dropout events [2]. These artifacts result in low biological fidelity, specifically a low signal-to-noise ratio (SNR), often submerging fine-grained biological signals within background noise[3]. Such noise confounds the identification of tissue microenvironments that govern cell fate and function. Therefore, a central computational challenge is the robust denoising and recovery of high-fidelity transcriptomic landscapes to enable the accurate characterization of spatial heterogeneity at biologically relevant scales [4].

To achieve high-fidelity reconstruction, current computational strategies typically integrate external multi-modal information, particularly high-resolution histology images, to guide expression prediction [5]. While incorporating such auxiliary context is theoretically beneficial, these multi-modal approaches generally rely on paired data and rigorous image registration [6]. This dependency limits their applicability in distorted tissues or scenarios where high-resolution auxiliary modalities are unavailable or misaligned [7]. To bypass these multi-modal dependencies, various uni-modal deep learning frameworks have been developed. Graph-based methods, such as GraphST [8] and STAGATE [9], leverage spatial topology to enhance representation learning; however, while effective for gene imputation and denoising at observed spots, their discrete nature fundamentally restricts them from performing spatial imputation or generating continuous expression landscapes at unmeasured coordinates. To address the need for continuous mapping, implicit neural representations (INRs [10]) have emerged to model continuous expression landscapes directly from spatial coordinates. Representative methods like STAGE and SUICA employ coordinate-based architectures to perform this mapping. However, STAGE relies on compressing high-dimensional data into a restrictive 2D latent space, creating an information bottleneck that risks discarding subtle biological variance [11]. Similarly, methods like SUICA incorporate spatial graphs by primarily utilizing nearest neighbor smoothing for denoising [12]. Such fixed-scale neighbor aggregation often fails to effectively capture complex microenvironments. Thus, these approaches struggle to perceive scale-dependent biological interactions, leaving the coordinate network functionally context-blind. This highlights the necessity for frameworks capable of explicitly encoding comprehensive, multi-scale contextual knowledge into coordinate representations, ensuring both local fidelity and robust microenvironmental awareness.

Here, we introduce the Cell Positioning System (CPS), a context-aware implicit neural representation framework designed to map physical coordinates to high-fidelity spatial transcriptomics via a privileged multi-scale context distillation strategy. To resolve the inherent context-blindness of standard INRs, CPS treats multi-scale tissue niches as privileged information, utilizing a framework where a teacher network explicitly models these niche structures during training to guide a student network in reproducing context-aware representations for graph-free inference [13]. Specifically, the teacher employs a multi-scale niche attention mechanism to adaptively integrate information from optimal receptive fields, thereby capturing effective biological interactions across varying neighborhood scales. By transferring this multi-scale contextual knowledge into the student network, CPS generates context-aware gene expression solely from spatial coordinates. This design effectively bypasses the dependency on auxiliary images and fixed-scale graph constraints, enabling the robust recovery of biologically meaningful signals even in data-sparse regions.

Comprehensive benchmarking on the human Dorsolateral Prefrontal Cortex (DLPFC) demonstrates that CPS achieves state-of-the-art performance in spatial and gene expression imputation and denoising tasks, exhibiting remarkable robustness against varying levels of data sparsity and noise. Leveraging its continuous generative nature, CPS further enables arbitrary-scale super-resolution, recovering high-resolution anatomical details that are invisible in raw resolution-limited data. Notably, the model offers intrinsic interpretability through its attention mechanism, which identifies the scale effective size (SES) of biological interactions, functioning as a computational lens to dissect tissue complexity. Finally, utilizing an efficient multi-scale graph tokenization strategy, CPS exhibits superior scalability, capable of processing large-scale datasets, such as Stereo-seq and Visium HD, with linear computational complexity. Moreover, extensive ablation studies validate the architectural design of CPS, confirming the indispensable contributions of each module to the overall predictive fidelity. By harmonizing continuous physical coordinates with discrete biological context, CPS provides a robust, interpretable, and scalable solution for resolving spatial heterogeneity.

## 2. Methods

### 2.1 Overview of CPS

The cell niche is a critical determinant of cell identity, driven by the complex interplay between spatial positioning and neighborhood context. To decode these interactions from physical coordinates and spatial gene expression co-correlations, we developed the cell positioning system (CPS), a graph-informed continuous field learning framework that integrates multi-scale niche context with physical coordinate field modeling to reconstruct high-fidelity transcriptomic profiles (Fig. 1). CPS adopts a multi-scale context-aware teacher network equipped with a graph tokenizer, which aggregates multi-hop neighbor gene expression features into spatial niche tokens. These tokens are dynamically integrated by a multi-scale niche attention mechanism. Crucially, this mechanism provides intrinsic interpretability by adaptively weighing local versus global dependencies, allowing CPS to identify the scale effective size (SES) of biological interactions that define distinct tissue structures. Parallel to this, a continuous field student network projects physical coordinates into high-dimensional frequency space via Fourier position mapping, modeling the tissue manifold using implicit neural representations (INR). Importantly, CPS employs a privileged information distillation (PID) strategy, guiding the student to internalize the multi-scale contextual knowledge encoded by the teacher, thereby harmonizing the modality discrepancy between discrete biological graphs and continuous spatial geometry. Optimized through a composite negative binomial (NB) and log-mean-squared error (LMSE) loss, CPS achieves superior fidelity in spatial and gene expression imputation and denoising. Furthermore, the INR-based generator enables arbitrary-scale super-resolution to recover high-resolution anatomical details, while an efficient training strategy based on multi-scale token pre-encoding ensures computational scalability for large-scale datasets, offering a unified solution for dissecting spatial heterogeneity.

**Figure 1.**
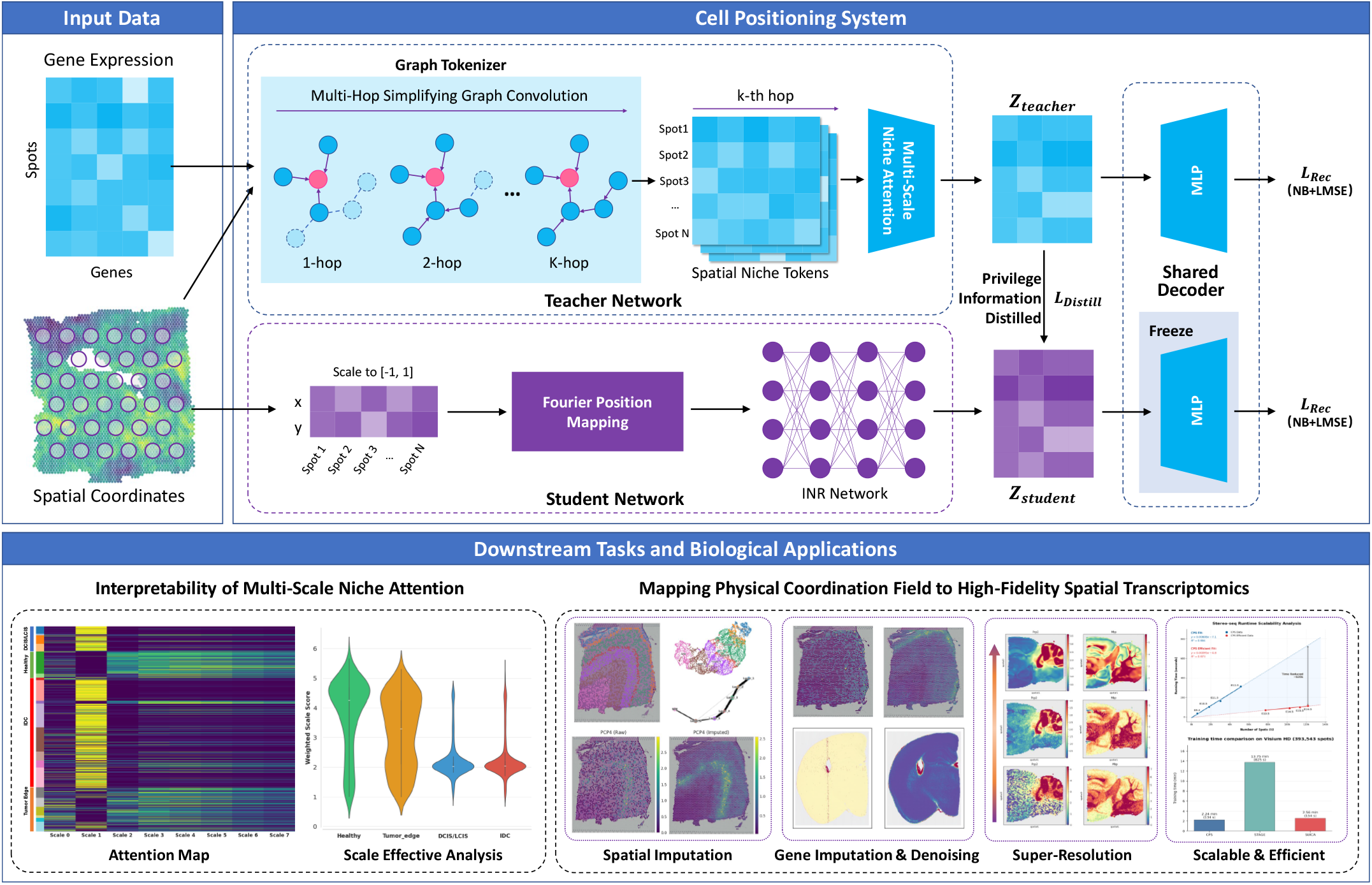
Overview of the Cell Positioning System (CPS) framework. **Input Data:** CPS utilizes raw gene expression matrices and the spatial coordinates of measured spots. **CPS Model Architecture**: The framework employs a teacher-student distillation paradigm. The **Teacher Network** (top) uses a Graph Tokenizer to aggregate multi-hop neighborhood features (from 1-hop to K-hop) with simplifying graph convolution into spatial niche tokens, which are integrated via a Multi-Scale Niche Attention mechanism to produce context-aware embeddings (*Z*_*teacher*_). The **Student Network** (bottom) projects physical coordinates into high-dimensional frequency space via Fourier Position Mapping, followed by an Implicit Neural Representation (INR) to generate coordinate-based embeddings (*Z*_*student*_). A Privileged Information Distillation (PID) strategy aligns the student with the teacher, while a Shared Decoder reconstructs expression profiles using a composite Negative Binomial (NB) and LMSE loss (*L*_*Rec*_). **Applications**: The unified framework supports downstream tasks including interpretability analysis (via attention weights), spatial imputation, gene imputation and denoising, and arbitrary-scale super-resolution.

### 2.2 Data description and preprocessing

We validated CPS across diverse spot-based spatial transcriptomics datasets from 10x Visium, Visium HD, and Stereo-seq platforms. The analysis included human dorsolateral prefrontal cortex (DLPFC) and human breast cancer (HBC), as well as mouse posterior brain section (MPBS), mouse brain (MBHD), and mouse developing embryos (MDE), demonstrating the framework’s adaptability to diverse tissue contexts and spatial resolutions. To derive robust feature representations, we processed raw gene expression profiles to select highly variable genes (HVGs) as model inputs, while explicitly preserving the original unnormalized count data for the output layer to calibrate the negative binomial reconstruction loss. We employ radius-based constraints for fixed-grid architectures and k-nearest neighbor connections for continuous coordinate-based platforms. Finally, physical coordinates were normalized to a unified range [-1, 1] to facilitate the stable convergence of the implicit neural representation. Comprehensive details regarding the data preprocessing pipelines and evaluation protocols (including specific masking strategies for gene and spatial imputation) are provided in the supplementary materials (Supplementary Note. 1 and 3).

### 2.3 Teacher network

The teacher network functions as a context-aware encoder designed to capture the multi-hop neighborhood context of the spatial niche. It consists of a graph tokenizer that generates multi-view representations of the local microenvironment, and a multi-scale niche attention module that adaptively fuses these representations.

#### 2.3.1 Graph tokenizer and multi-hop simplifying graph convolution

To efficiently capture multi-scale structural information across varying spatial ranges, we employ a parallel simplifying graph convolution network [14]. Given the gene expression matrix **X** and the symmetrically normalized adjacency matrix **Ã**, input features are first projected to a latent space **H**^(0)^ via a linear layer with GELU activation. We then generate spatial niche tokens by propagating these latent features over a set of neighborhood scales κ = {*k*_1_, …, *k*_*S*_}:

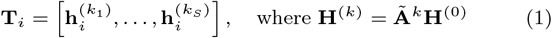

Here, **T**_*i*_ ∈ ℝ^*S*×*d*^ represents the multi-view neighborhood context for spot *i*, where 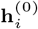 denotes the local identity and 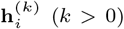 aggregates information from the *k*-hop neighborhood.

#### 2.3.2 Multi-scale niche attention

To effectively integrate these multi-scale spatial niche tokens, we propose a specialized attention mechanism where the spot’s local state queries its surrounding niches. We first add learnable scale position embeddings to **T** to preserve scale identity. For each spot *i*, the Query (**q**_*i*_) is derived exclusively from its local 0-hop token, while Keys (**K**_*i*_) and Values (**V**_*i*_) are projected from the full token sequence. The context-aware spot representation **c**_*i*_ is computed as a weighted sum:

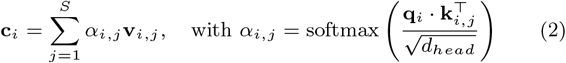

The attention weights *α*_*i,j*_ reflect the importance of specific spatial scales. The final output **Z**_*teacher*_ is obtained by passing **c**_*i*_ through a linear layer and adding a residual connection from the local features **H**^(0)^ to preserve intrinsic gene expression signals.

#### 2.3.3 Entropy regularization

Standard attention mechanisms typically initialize near a uniform distribution, which risks trapping the model in an over-smoothed state with vanishing gradients during early training. To break this initial symmetry and prevent the attention weights from getting stuck in a uniform distribution, we introduce an entropy regularization term [15]. We compute the entropy ℋ of the average attention weights and penalize deviations from a uniform distribution prior using a hinge loss:

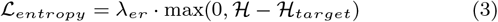

where ℋ_*target*_ = 0.5 ln(*S*). By actively forcing the entropy down from its initial maximum, this regularization ensures the model quickly escapes the uniform trap, actively exploring and committing to discriminative spatial scales rather than collapsing into a context-blind average.

### 2.4 Student network

Parallel to the context-aware teacher network, the student network functions as a coordinate-based generator designed to model the tissue as a continuous field. By mapping physical coordinates directly to latent representations, this module enables arbitrary-scale continuous inference and captures the continuous expression landscape of the tissue.

Standard multi-layer perceptrons (MLPs) suffer from spectral bias, rendering them inefficient at learning high-frequency functions such as sharp tissue boundaries. To overcome this, we employ implicit neural representations (INRs) augmented with a Gaussian Fourier feature mapping [16]. Given normalized spatial coordinates **x** ∈ ℝ^2^, we project them into a high-dimensional frequency space before feeding them into the network. The mapping function *γ*(·) is defined as:

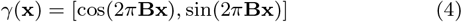

where **B** ∈ ℝ^*m*×2^ is a Gaussian random matrix sampled from 𝒩 (0, *σ*^2^), and *m* denotes the number of frequency bands. The hyperparameter *σ* controls the frequency spectrum; a larger *σ* enables the model to capture finer spatial details. The encoded features *γ*(**x**) are then processed by an MLP with SiLU activations and LayerNorm to generate the coordinate-based latent representation **Z**_*student*_.

### 2.5 Generative reconstruction

To reconstruct high-fidelity gene expression, we employ a shared decoder that maps latent embeddings from both teacher and student networks to the transcriptomic manifold. The decoder predicts the mean expression ***µ***_*i*_ for spot *i* by scaling the softmax-normalized logits with the observed library size *l*_*i*_:

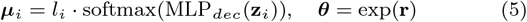

where ***θ*** denotes the learnable gene-specific dispersion.

To strictly model the sparse and over-dispersed count data, we adopt the negative binomial (NB) distribution [17]. The likelihood of observing gene counts **x**_*i*_ is given by:

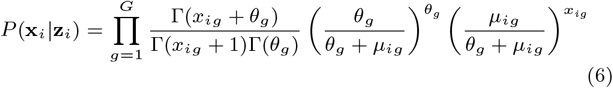

We minimize the negative log-likelihood (NLL) as the primary objective. To further stabilize training, we incorporate an auxiliary log-mean-squared error (LMSE) term. The total reconstruction loss ℒ_*rec*_ is thus formulated as:

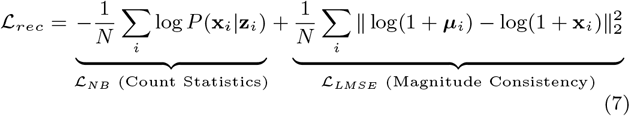

### 2.6 Asymmetric co-training strategy

To effectively harmonize discrete spatial graph and continuous spatial geometry, we propose an asymmetric co-training strategy.

#### Phase 1: context-guided manifold construction

In the first phase of each iteration, we update the parameters of the teacher network and the shared decoder. The objective is to minimize the reconstruction loss and entropy regularization:

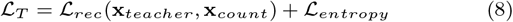

Crucially, the student network is paused during this step. This phase constructs a high-fidelity, context-aware latent manifold that serves as the stable biological anchor for the subsequent alignment.

#### Phase 2: coordination-driven alignment via privileged information distillation

A key challenge in spatial transcriptomics is that spatial coordinates alone often lack sufficient information to infer complex spatial states defined by local interactions. To bridge this gap, we implement a privileged information distillation (PID) strategy during the second phase. Here, the local neighborhood context which accessible only to the teacher via the graph is treated as privileged information. We update only the student network to align with the teacher’s latent space while simultaneously minimizing reconstruction error. The student optimization objective is defined as:

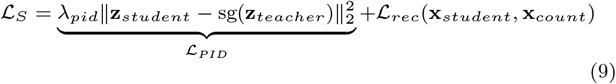

where sg(·) denotes the stop-gradient operator. The ℒ_*PID*_ term forces the student to internalize the structural knowledge captured by the teacher, effectively inferring the neighborhood structure solely from spatial coordinates.

Crucially, the shared decoder is frozen during this phase. By decoupling the optimization processes, this asymmetric two-stage design safely avoids gradient conflicts within the coupled computational graph. Furthermore, keeping the decoder frozen anchors the student’s latent space to the exact biological manifold learned by the teacher. This constraint forces the student to map physical coordinates strictly into the existing high-frequency representations, guaranteeing that the coordinate-based network genuinely captures complex microenvironmental contexts rather than simply collapsing into a smoothed, context-blind approximation.

## 3. Results

### 3.1 CPS achieves the best performance in spatial and gene imputation on the DLPFC dataset

To rigorously evaluate the performance of CPS in recovering biologically meaningful signals from sparse spatial transcriptomics data, we benchmarked our framework against STAGE, SUICA, and two representative graph-based methods, STAGATE and GraphST, using the human dorsolateral prefrontal cortex (DLPFC) dataset. The evaluation encompassed both gene imputation (recovering dropout values in observed spots) and spatial imputation (predicting expression in unmeasured spatial locations). Notably, because the inherent discrete nature of graph-based algorithms requires pre-constructed spatial adjacency matrices, STAGATE and GraphST are structurally restricted to *in situ* denoising and were therefore only evaluated in the gene imputation task.

We first quantified the reconstruction accuracy using five metrics: mean absolute error (MAE), mean squared error (MSE), and Pearson correlation coefficient (PCC) (Fig. 2A), alongside cosine similarity and Spearman correlation coefficient (SCC) (Supplementary Fig. S18). In the gene imputation task, CPS demonstrated robust superiority, consistently achieving the lowest MAE and MSE, as well as the highest Pearson correlation and Cosine similarity compared to all four baselines. While STAGE exhibited a slightly higher Spearman correlation when evaluated globally across all spot expressions, CPS reclaimed the lead when specifically evaluating non-zero expression values (Supplementary Fig. S18). This distinction is crucial: it suggests that while baseline methods may effectively replicate background sparsity, CPS is significantly more accurate at ranking and recovering relative expression levels within biologically active regions.

**Figure 2.**
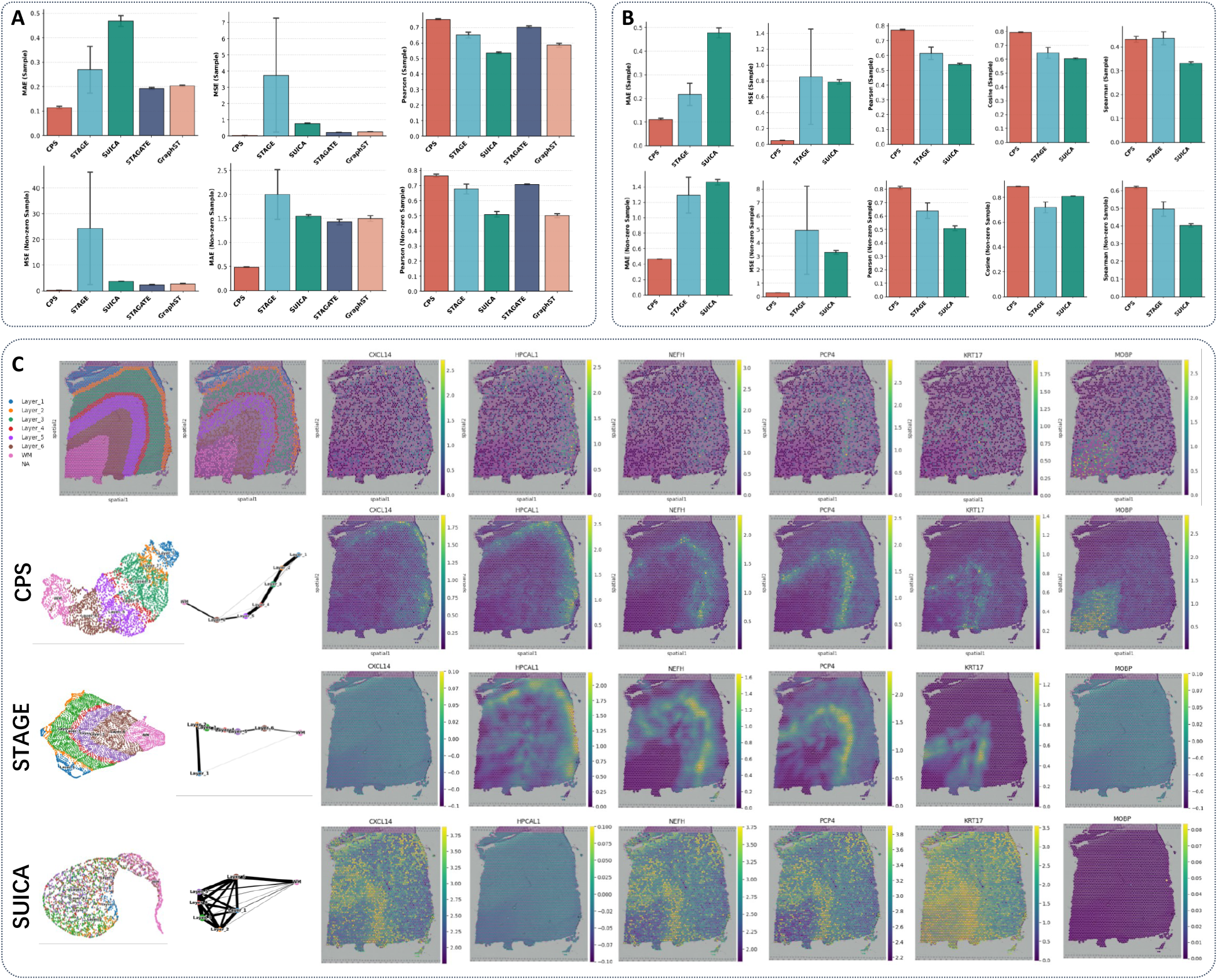
Benchmarking CPS against state-of-the-art methods on the DLPFC dataset. A. Quantitative comparison of gene imputation performance using MAE, MSE, and Pearson metrics across CPS, STAGE, SUICA, STAGATE, and GraphST. B. Quantitative comparison of spatial imputation performance on held-out test spots for CPS, STAGE, and SUICA. C. Visualization of spatial imputation results. Left: PAGA trajectories and UMAP embeddings colored by ground-truth layers, showing the structural reconstruction quality. Right: Spatial expression maps of representative layer-marker genes (*CXCL14, HPCAL1, NEFH, PCP4, KRT17, MOBP*). CPS recovers distinct laminar patterns with higher fidelity compared to STAGE and SUICA.

In the spatial imputation task (Fig. 2B), where the model must generalize to completely unseen spatial coordinates, CPS maintained its decisive lead over STAGE and SUICA. It achieved a substantial reduction in MSE and surpassed both competitors across all correlation metrics, demonstrating that our continuous coordinate-based generator captures the underlying spatial expression landscape much more effectively than restricted latent bottlenecks or fixed-scale interpolation strategies. Beyond quantitative metrics, we assessed the model’s ability to reconstruct the distinct laminar architecture of the DLPFC. We visualized the latent space representations and PAGA trajectories [18] derived from the spatially imputed data (Fig. 2C). The PAGA graph generated by CPS revealed a clean, quasi-linear trajectory that perfectly mirrored the biological transition from Layer 1 through Layer 6 to the White Matter (WM). The nodes were well-separated yet connected in a biologically logical sequence. In contrast, the SUICA-derived trajectory appeared densely interconnected and disordered, failing to distinguish the sequential laminar progression. Visual inspection of imputed gene expression maps further highlighted the advantages of CPS (Fig. 2C). Focusing on known layer-specific markers such as *CXCL14* (Layer 2), *HPCAL1* (Layer 2/3), *NEFH* (Layer 3-5), *PCP4* (Layer 5), and *MOBP* (White Matter), CPS generated crisp expression domains with well-defined edges. Conversely, SUICA tended to over-smooth expression patterns, resulting in blurred layer boundaries, while STAGE reconstructions often contained spotty artifacts. Finally, evaluating gene imputation capabilities at the single-spot level (Supplementary Fig. S2) confirmed that CPS intelligently recovers missing information based on learned spatial-molecular dependencies, transforming severely dropped-out raw data into highly distinct, spatially coherent layer-specific enrichments.

### 3.2 CPS enables continuous super-resolution reconstruction of mouse posterior brain architectures

We next applied CPS to the mouse posterior brain dataset to evaluate its capacity for super-resolution (SR) reconstruction, enabling the visualization of high-resolution anatomical details beyond the physical resolution limit of the Visium platform. To validate the fidelity of the reconstructed patterns, we compared the model outputs against the histological reference (H&E staining, Fig. 3A), focusing on complex tissue architectures. We first quantified the quality of the generated expression fields using three complementary metrics: Moran’s I [19], Robust Contrast-to-Noise Ratio (CNR [20]), and Geary’s C (Fig. 3B). Moran’s I measures global spatial autocorrelation, serving as a proxy for the overall spatial continuity of gene expression, while Robust CNR evaluates the distinguishability of biological signals from background noise. Crucially, to explicitly confirm that CPS avoids artificial mathematical over-smoothing, we introduced Geary’s C, which is highly sensitive to localized spatial structural transitions. While simple smoothing acts as a low-pass filter that artificially washes out local differences, a lower Geary’s C score indicates the successful preservation of high-frequency spatial features and sharp boundary contrast. Across a panel of representative genes (*Pcp2, Mbp, Hpca, Calb1, Gabra6, Cux2* and *Car3*), the raw Visium data exhibited relatively poor scores due to inherent sparsity and dropout events. In contrast, the CPS-reconstructed data at the original resolution (CPS SR-X1) showed a dramatic improvement across all metrics. Notably, this high fidelity was maintained consistently across super-resolution scales (X2, X4, and X6). The model maintained high Moran’s I and CNR while preserving low Geary’s C scores. This indicates that its inference at sub-spot resolutions does not introduce noise or blurred artifacts, but rather accurately preserves the underlying biological structures. Visual comparison of gene expression maps within highly structured regions of interest (ROIs) further confirms these quantitative findings (Fig. 3C and Supplementary Fig. S16). While raw data suffers from pixelated noise and standard k-NN smoothing generates biologically inaccurate blurred boundaries, CPS effectively denoises while restoring crisp continuous expression domains. Progressively upsampling the implicit neural field to higher resolutions (X2, X4, and X6) revealed intricate structural details that tightly align with true anatomical features in the corresponding H&E images. For instance, across a panel of representative marker genes (*Calb1, Hpca* and *Cux2*), CPS sharply delineates specific tissue domains and layers with smooth yet highly contrasted boundaries. These targeted ROI analyses visually and mathematically prove that CPS achieves true biological super-resolution rather than mere mathematical smoothing.

**Figure 3.**
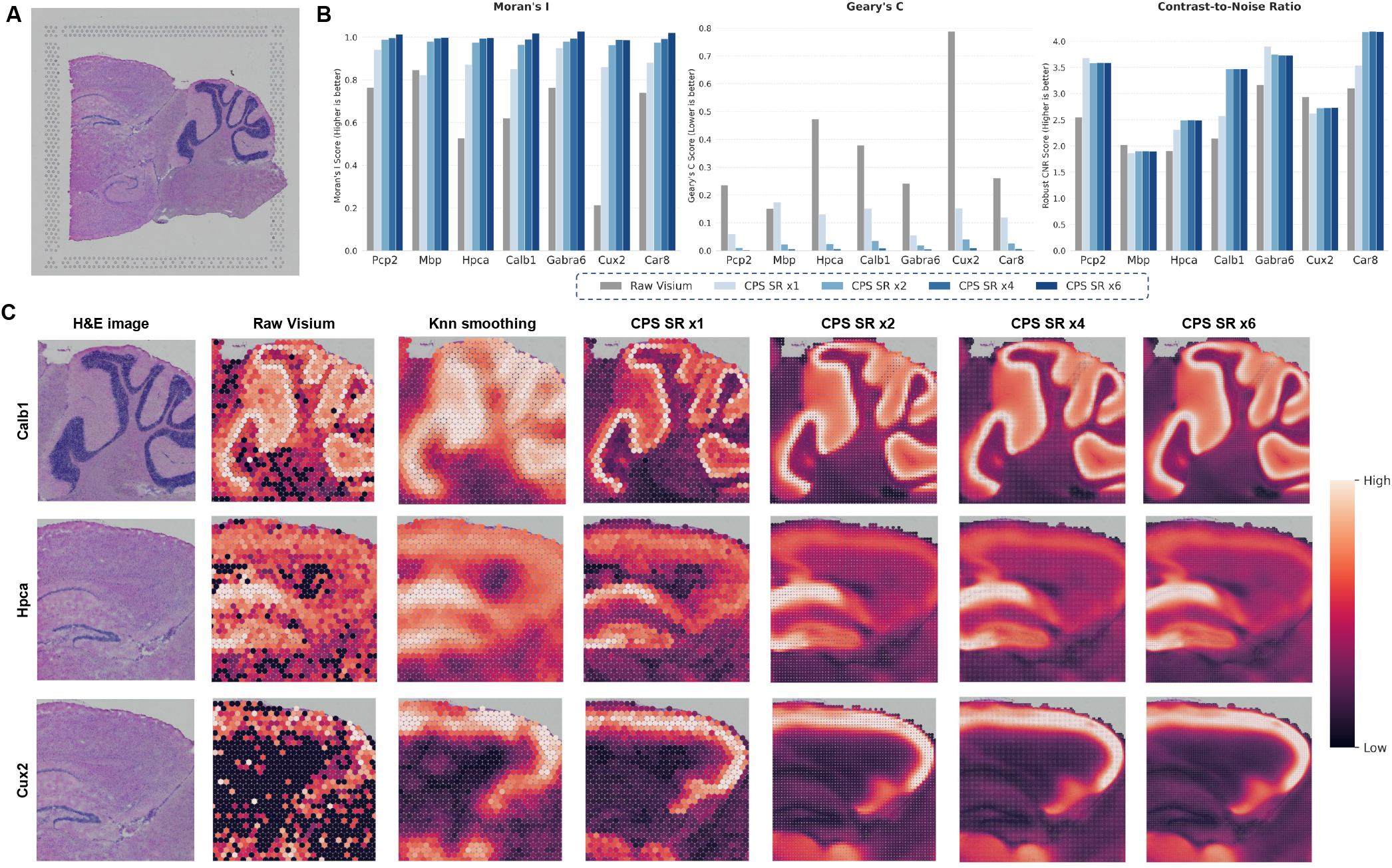
Continuous super-resolution reconstruction via CPS. **A**. H&E stained histological reference of the mouse posterior brain. **B**. Quantitative comparison of Moran’s I, Geary’s C, and Robust Contrast-to-Noise Ratio (CNR) metrics across resolutions (*×*1 to *×*6). **C**. Multi-scale visualization of representative marker genes (*Calb1, Hpca, Cux2*) within structural Regions of Interest (ROIs). Columns display the corresponding H&E image, raw data, k-NN smoothing, CPS denoised output (CPS SR *×*1), and continuous super-resolution inferences (*×*2, *×*4, *×*6). CPS preserves sharp boundary contrast compared to k-NN smoothing, accurately recovering fine anatomical details obscured in raw data.

### 3.3 CPS decodes microenvironmental heterogeneity with interpretable multi-scale niche attention patterns

To validate the interpretability of CPS, we analyzed a human breast cancer dataset characterized by a complex tissue architecture comprising healthy tissue, tumor edges, and distinct lesion regions (DCIS/LCIS and IDC) (Fig. 4A). The model learned a dynamic attention distribution across 8 spatial scales (Fig. 4D). Quantitative analysis of the attention weights revealed a context-dependent behavior where lesion regions (DCIS/LCIS and IDC) relied predominantly on local features and the tumor edge consistently recruited broader context information (Figs. 4B-C). Scale effective size (SES) and scale entropy (SE) metrics further confirmed this trend and demonstrated that the tumor edge exhibits higher receptive field complexity compared to the lesion cores (Figs. 4E-F).

**Figure 4.**
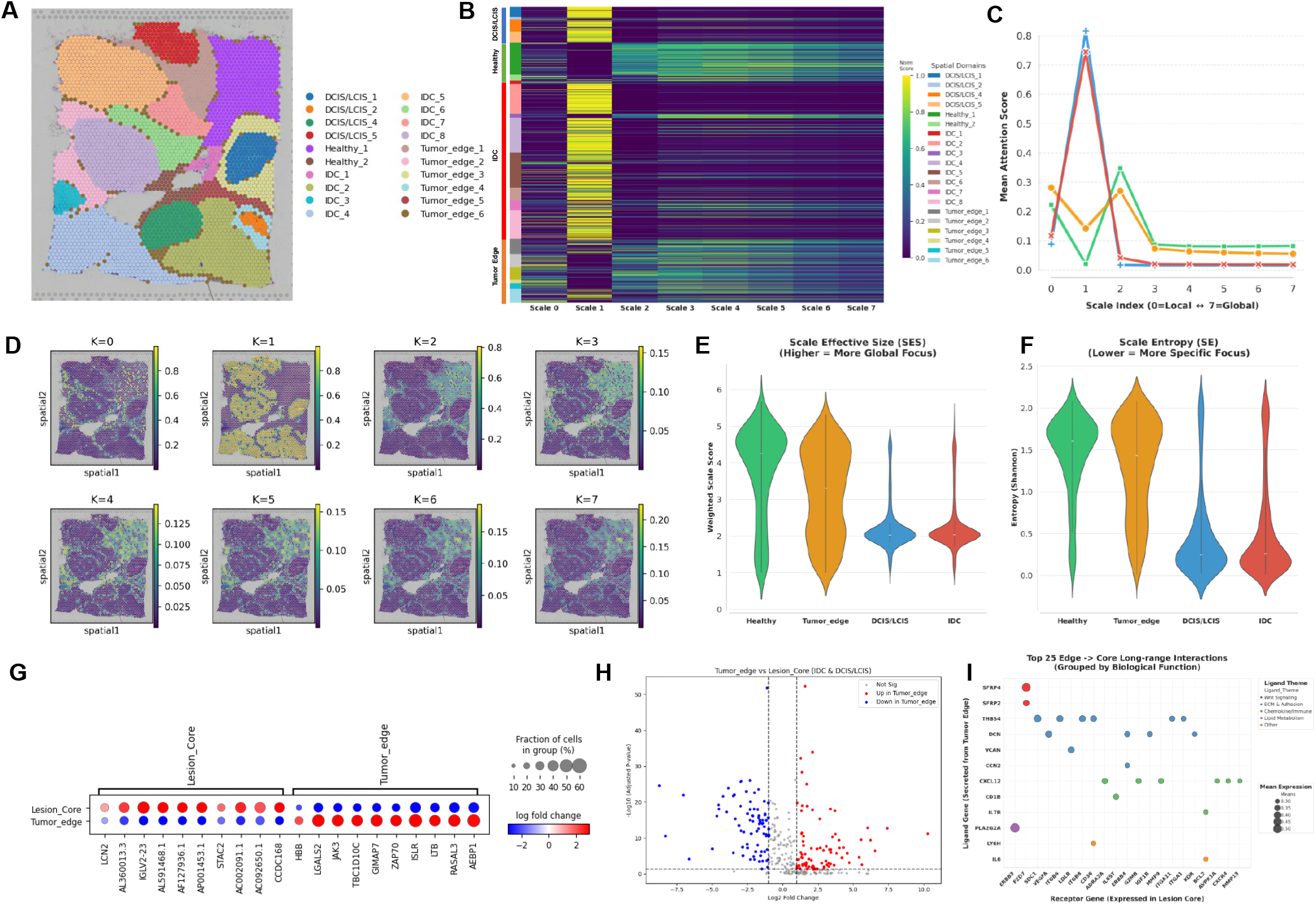
Interpretable multi-scale niche attention of CPS. **A**. The ground truth of the Human Breast Cancer dataset. **B**. Column-normalized heatmap of attention scores across spatial domains. **C**. Line plot of average attention scores across scales for four tissue domains. **D**. Visualization of attention scores across 8 scales (ranging from *K* = 0 to *K* = 7). **E**. Violin plot of scale effective size (SES) on four tissue domains. **F**. Violin plot of scale entropy (SE) on four tissue domains. **G**. Dotplot of differentially expressed genes (DEGs) between the Lesion Core (IDC and DCIS/LCIS) and Tumor Edge, computed using CPS imputed data. **H**. Volcano plot of the differential expression analysis (Tumor Edge vs. Lesion Core) using CPS output. **I**. Ligand-receptor interaction analysis based on CPS derived DEGs, mapping the spatial cell-cell communication active at the tumor edge.

We next investigated the molecular basis driving the model’s multi-scale niche attention and demonstrated the downstream utility of CPS. We grouped the structural domains into Lesion Core (IDC and DCIS/LCIS) and Tumor Edge and performed differential expression analysis (DEA). Utilizing the raw Visium data for DEA yielded highly limited results (Supplementary Fig. S17) because severe dropout and high sparsity compromised the statistical power. This resulted in very few significant genes and provided limited spatial insights. Conversely, performing DEA on the reconstructed expression profiles of CPS recovered the biological signals (Figs. 4G-H). CPS enhanced the statistical power and uncovered a broader set of differentially expressed genes (DEGs) that exhibit distinct spatial localization at the tumor edge.

To determine whether the expanded SES observed at the tumor edge relates to underlying biological interactions rather than spatial optimization artifacts, we subjected these derived DEGs to a ligand-receptor cell communication analysis using Squidpy [19] (Fig. 4I). The analysis identified multiple active spatial interaction pairings at the tumor boundary. Prominent receptor-ligand pairs enriched in the high-SES Tumor Edge (such as *THBS4* -*ITGB6* and *DCN* - *VEGFA*) are classically associated with extracellular matrix (ECM) remodeling, angiogenesis, and tumor-stroma crosstalk. This suggests that the broader receptive field learned by CPS corresponds to regions with active intercellular communications, demonstrating its utility in characterizing complex spatial microenvironments.

### 3.4 Ablation studies, robustness, and scalability of the CPS framework

To evaluate the architectural design of CPS, we conducted systematic ablation studies by removing or replacing key components (Supplementary Figs. S8-S13). Omitting the privileged information distillation (PID) loss or the asymmetric co-training strategy severely compromised the model’s ability to recover sharp spatial boundaries. This empirically confirms that decoupling the optimization processes prevents gradient conflicts and ensures the coordinate-based student network captures complex spatial contexts rather than collapsing into a smoothed approximation. Similarly, removing the Fourier feature mapping resulted in a loss of high-frequency anatomical details, while altering the composite NB+LMSE objective reduced reconstruction fidelity on over-dispersed count data. We also assessed the parameter sensitivity and robustness of CPS (Supplementary Figs. S5-S7 and S14-S15). The evaluations confirmed that the framework maintains high generative performance across a wide range of hyperparameter settings. Furthermore, under simulated conditions of extreme data sparsity and elevated noise, CPS consistently maintained high reconstruction quality and outperformed baseline methods. Finally, we evaluated the framework’s scalability on large-scale datasets from Visium HD and Stereo-seq platforms. CPS employs an efficient training mode based on multi-scale token pre-encoding to decouple graph aggregation from the iterative training loop. Runtime profiling (Supplementary Fig. S19) demonstrated that this approach achieves linear complexity with respect to the number of spots. Specifically, CPS achieved rapid convergence on a dense Visium HD dataset comprising over 390,000 spots in just 2.24 minutes. This demonstrates its practical scalability for next-generation spatial transcriptomics applications.

## 4. Conclusion

In this study, we presented the Cell Positioning System (CPS), a coordinate-based continuous generative framework designed to reconstruct high-fidelity spatial transcriptomic landscapes from sparse and noisy measurements. By employing a privileged multi-scale context distillation strategy, CPS effectively harmonizes the continuous nature of physical geometry with the discrete structural context of biological microenvironments. This methodological innovation resolves the inherent context-blindness of coordinate-based representations without relying on auxiliary histology images or rigid graph constraints. Our comprehensive benchmarking demonstrates that CPS not only achieves state-of-the-art performance in spatial and gene imputation and denoising but also enables arbitrary-scale super-resolution to uncover high-resolution anatomical details. Uniquely, the multi-scale niche attention mechanism serves as an interpretable method, quantifying the scale effective size (SES) of biological interactions to decode complex microenvironments. Coupled with linear computational scalability, CPS offers a robust and efficient solution for next-generation, high-density spatial transcriptomics. Future work will extend this framework to three-dimensional tissue reconstruction and multi-modal integration, further advancing our ability to dissect spatial heterogeneity in complex biological systems.

## Data availability

The datasets analyzed in this study are publicly available from their respective repositories:

- The human Dorsolateral Prefrontal Cortex (DLPFC) dataset is available via the *spatialLIBD* package or at http://research.libd.org/spatialLIBD/.
- The Human Breast Cancer, Mouse Posterior Brain (Visium) and Mouse Brain Visium HD datasets can be downloaded from the 10x Genomics website (https://www.10xgenomics.com/resources/datasets).
- The Stereo-seq Mouse Embryo Atlas is accessible through the MOSTA database (https://db.cngb.org/stomics/mosta/).

The source code for CPS, including model implementation, training scripts, and tutorials, is freely available on GitHub at https://github.com/tju-zl/CPS. All pre-processed data and scripts required to reproduce the benchmarking results and figures are also provided in the GitHub repository.

## Author contributions statement

L.Z., K.C. and L.W. conceived the experiments, L.Z. and S.Z. conducted the experiments, L.Z., K.C. and L.W. analysed the results. L.Z., K.C., S.Z., S.L. and L.W. wrote and reviewed the manuscript.

