## Supplementary for "CPS: Mapping Physical Coordinates to High-Fidelity Spatial Transcriptomics via Privileged Multi-Scale Context Distillation"

**Abstract**

**Motivation:** Spatial transcriptomics enables the dissection of tissue heterogeneity within native contexts, yet current platforms are inherently constrained by high sparsity and low signal-to-noise ratios that obscure fine-grained biological signals. Current efforts to recover these signals are limited by image registration dependencies or the inherent context-blindness of implicit neural representations.

**Results:** We introduce the Cell Positioning System (CPS), a context-aware implicit neural representation framework designed to map physical coordinates to high-fidelity spatial transcriptomics via a privileged multi-scale context distillation strategy. CPS treats multi-scale tissue niches as privileged information, employing a teacher network equipped with a multi-scale niche attention mechanism to capture adaptive biological interactions during training. This structural knowledge is explicitly distilled into a student coordinate network, enabling the generation of context-aware expression landscapes solely from spatial coordinates during inference. Benchmarking on the DLPFC dataset demonstrates that CPS achieves state-of-the-art performance in spatial and gene expression imputation and denoising. Furthermore, CPS enables super-resolution to recover high-resolution mouse brain anatomical details and offers interpretability by identifying the scale effective size of biological interactions within human breast cancer tissues. Finally, the framework exhibits superior scalability for large-scale datasets with linear computational complexity.

**Availability:** Software is available online athttps://github.com/tju-zl/CPS.

**This file contains the following materials:**

1. Supplementary Notes 1-4
2. Supplementary Figures 1-19
3. Supplementary Tables 1

### Supplementary Notes

#### Supplementary Note 1. Detailed data preprocessing pipelines

To derive informative feature representations from the raw data, we processed the gene expression profiles using the Scanpy framework. Initially, genes expressed in fewer than three spots were filtered out to reduce noise. The data were then normalized by library size to a fixed total count, log-transformed, and scaled to zero mean and unit variance. Based on dispersion metrics, the top 3,000 highly variable genes (HVGs) were selected as the input for CPS network. Crucially, distinct from these transformed inputs, we retained the raw, unnormalized integer counts for the final output layer to compute the negative binomial (NB) and log mean square error (LMSE) loss, ensuring the model learns the underlying count distribution directly from observed data. To capture the local neighborhood graph intrinsic to each platform, we constructed spatial adjacency graphs tailored to the specific data structures. For platforms characterized by fixed grid architectures, such as 10x Visium, we employed a radius-based nearest neighbor (RKNN) graph, where the cutoff radius was determined based on the theoretical center-to-center distance between spots. Conversely, for datasets with irregular cell distributions like Stereo-seq, we utilized a k-nearest neighbor (KNN) graph based on Euclidean distances to ensure robust connectivity across varying densities. Finally, to harmonize the spatial and genomic modalities and facilitate the convergence of the Fourier position mapping, all physical coordinates were linearly scaled to the range [-1, 1].

For more implementation details of preprocessing, please refer to our reproducibility notebook on: https://github.com/tju-zl/CPS/.

#### Supplementary Note 2. Definition of scale effective size (SES) and scale entropy (SE) in multi-scale niche attention patterns

To strictly quantify the model's effective receptive field, we analyze the attention weights $\alpha_{i,k}$ learned by the Teacher network across $K$ spatial scales. We first compute the normalized scale probability distribution$p_{i,k} \propto mean(\alpha_{i,k})$. The SES and SE are then defined as the expectation and entropy of this distribution, respectively:

$$\mathrm{SES}_i = \sum_{k=0}^{K-1} (k+1) \cdot p_{i,k}, \qquad \mathrm{SE}_i = -\sum_{k=0}^{K-1} p_{i,k} \log p_{i,k}$$

where $k \in \{0, 1, .., K-1\}$ denotes the neighborhood hop index. SES proxies the expected radius of the receptive field (ranging from 1 to K), while SE measures the diversity of spatial integration, with higher entropy indicating reliance on multi-scale contexts.

#### Supplementary Note 3. Baseline Implementation Details and Evaluation Protocols

To rigorously evaluate the performance of CPS against state-of-the-art methods, we conducted a comprehensive benchmark using the DLPFC dataset (12 slices) under a strict 50% down-sampling setting for both spatial imputation and gene imputation tasks.

For the spatial imputation task, we compared CPS against STAGE and SUICA. We implemented the STAGE method strictly adhering to its official tutorial. For the SUICA method, we identified potential data leakage risks inherent in its original two-step sampling strategy. To mitigate this, we synchronized the training and testing indices between its Graph Neural Network (GNN) and Fourier Feature Network (FFN) modules and fixed random seeds to ensure reproducibility. We acknowledge that its training framework may still implicitly utilize test set information for hyperparameter optimization in SUICA. Furthermore, while the original SUICA documentation implies ambiguity regarding input transformation, we observed training failures in the absence of logarithmic scaling. We consequently applied count normalization alongside necessary transformations to ensure model convergence.

For the gene imputation task, we expanded the baseline comparisons to include two representative graph-based methods, STAGATE and GraphST. Because the inherent discrete nature of these algorithms requires pre-constructed spatial adjacency matrices, they are structurally restricted to in situ denoising and cannot perform continuous spatial imputation on unmeasured coordinates. Since none of the baseline methods provided specific implementation guidelines for gene imputation, we adopted an external validation framework analogous to CPS. We manually masked the input data and preserved the corresponding ground truth values prior to model input. STAGATE and GraphST were executed using their default hyperparameter settings as recommended in their official tutorials.

Quantitative evaluation was performed using mean squared error (MSE), mean absolute error (MAE), cosine similarity (CS), Pearson correlation coefficient (PCC), and Spearman correlation coefficient (SCC). To ensure fairness in calculating metrics for count-based data, we applied a Log(x+1) transformation to both the predicted values and the ground truth prior to computation.

These metrics were assessed under two distinct conditions across all features and restricted to non-zero features to provide a robust evaluation of recovery performance. The final reported metrics represent the average values across all 12 DLPFC slices accompanied by their respective standard deviations. Beyond statistical metrics, we further validated the biological relevance of the imputed data by examining layer-specific marker genes in the DLPFC and analyzing the spatial architecture of the generated omics data through UMAP projections and PAGA trajectory inference.

#### Supplementary Note 4. Computational Cost and Hardware Requirements

To provide a transparent evaluation of the computational cost and hardware requirements of the CPS, all experiments were conducted on a workstation equipped with an Intel Core i9-12900K processor, 64 GB of system RAM, and a single NVIDIA GeForce RTX 3090 GPU with 24GB of VRAM, operating within an Ubuntu 20.04 environment via Windows Subsystem for Linux 2 (WSL2). For our batching strategy in the efficient mode (only for large dataset over 50k spots of single slice), we structured the input data into mini-batch of size 256 in the efficient mode. Regarding memory usage, the peak VRAM consumption during the training phase reached approximately 11686 MB. Meanwhile, the data preprocessing and tokenization stages were strictly CPU-bound, consuming up to 11.9 GB of the system RAM. Under these hardware specifications and configurations, the average training runtime was 5.56s per epoch, and a complete training pass for a standard dataset containing 121767 cells required only 98 seconds. This full accounting explicitly demonstrates that the proposed method remains computationally tractable and scales efficiently to large-scale datasets.

### Supplementary Figures

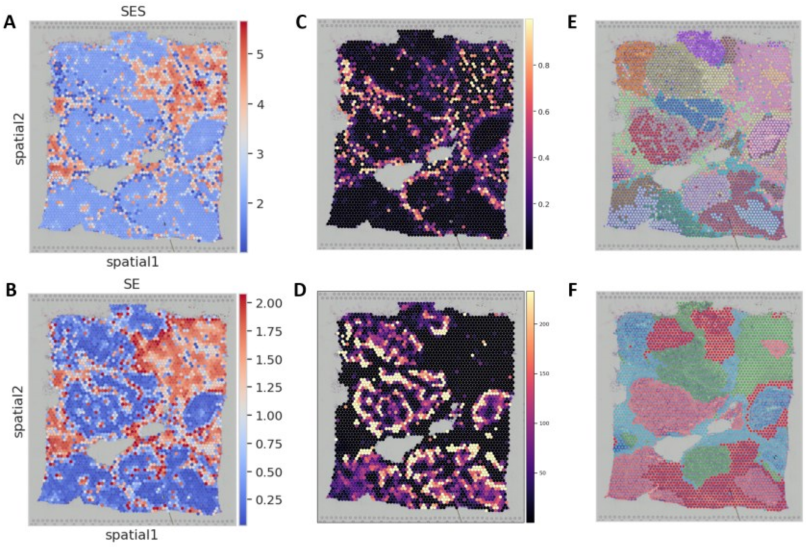


**Supplementary Figure 1.** A. The visualization of scale effective size of HBC dataset. B. Scale Entropy visualization. C. The scale 0 visualization. D. The enhanced first 2 scales visualization (the first two scales are divided by the remaining scales). E. The clustering labels of teacher latent space. F. The clustering labels of student latent space.

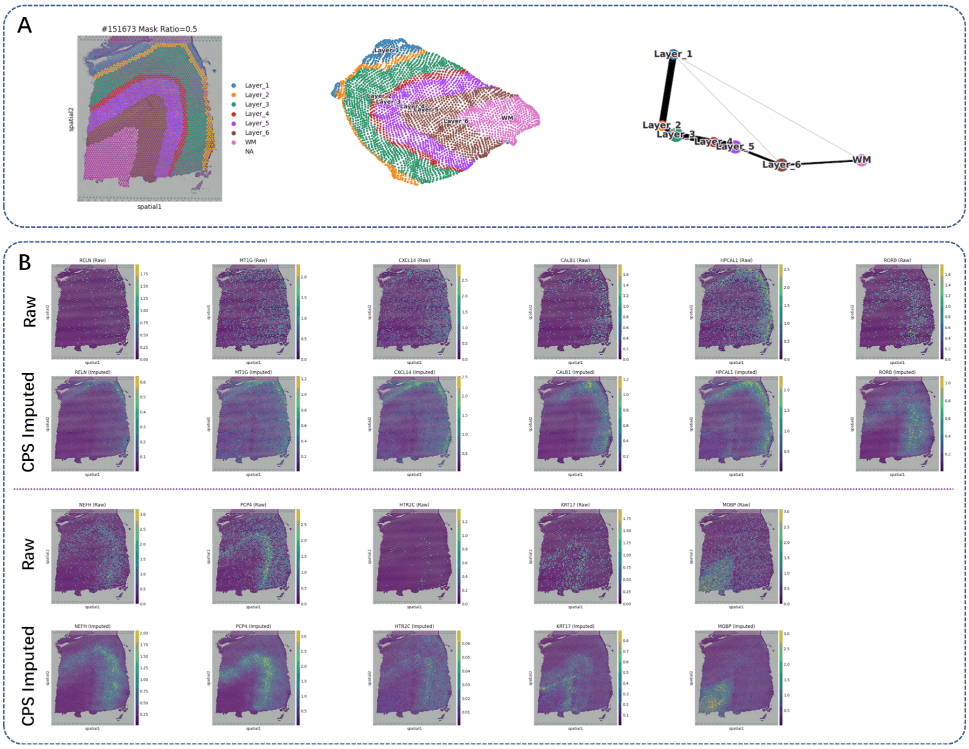


**Supplementary Figure 2.** Visualization of Gene Imputation Performance. Comparison between Raw expression and CPS imputed expression for representative genes. CPS effectively recovers continuous spatial patterns from sparse input data.

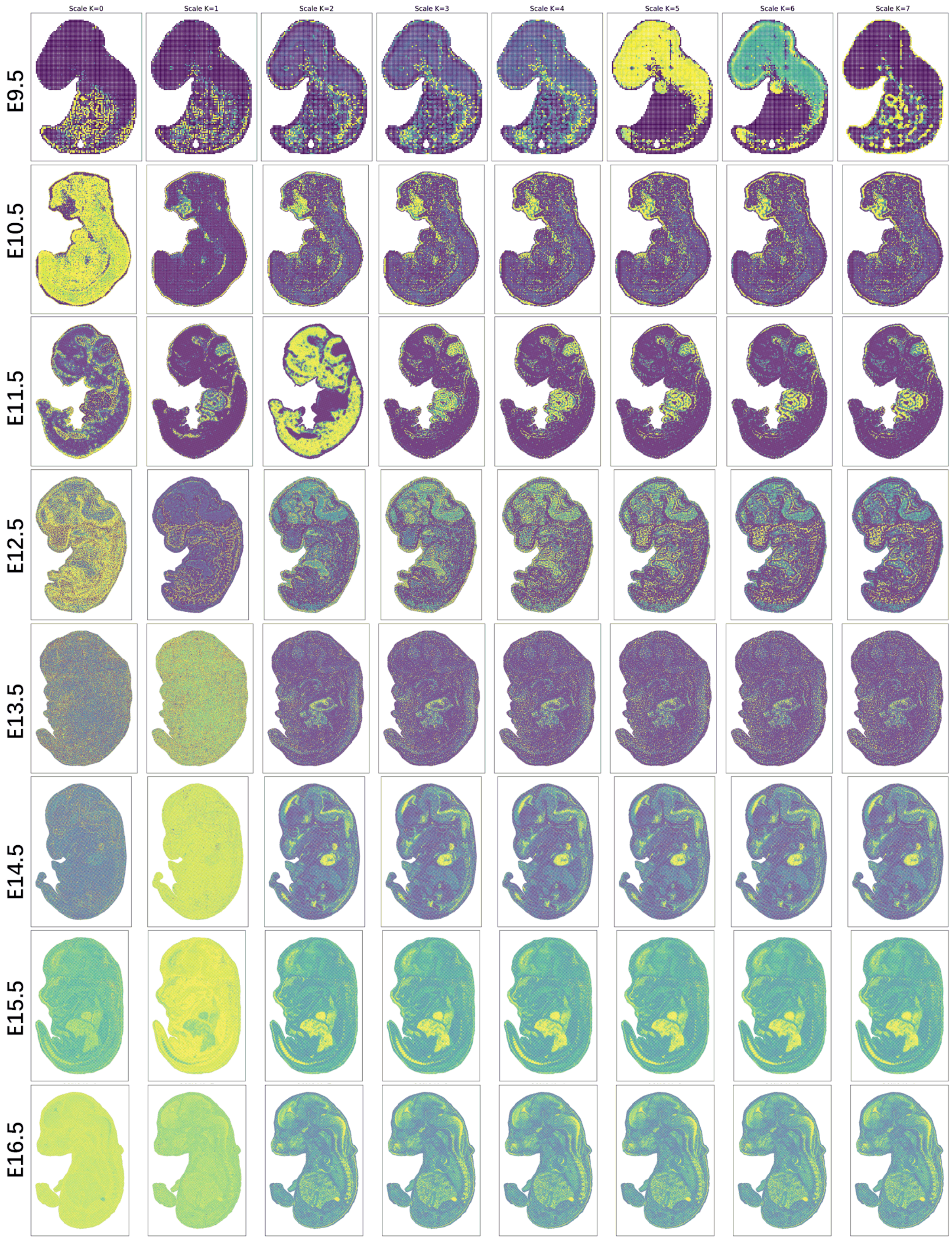


**Supplementary Figure 3.** Multi-scale niche attention patterns on Stereo-seq mouse developing embryo dataset.

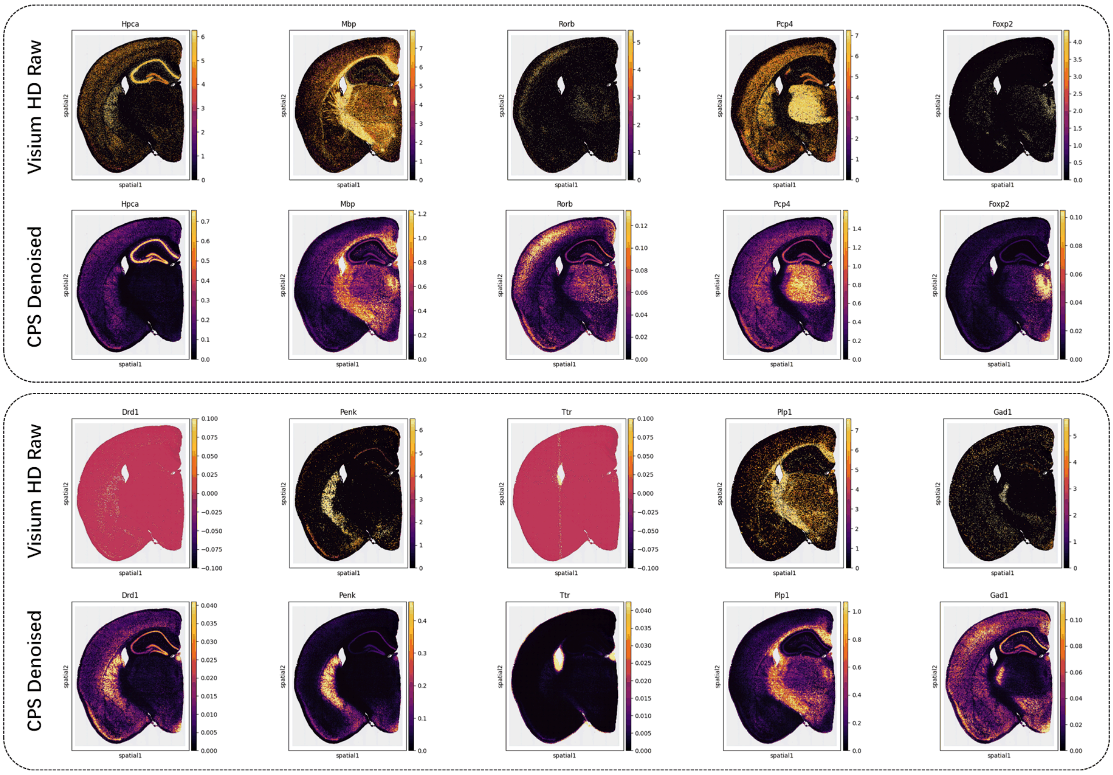


**Supplementary Figure 4.** Gene expression denoising of CPS on 10x Visium HD mouse brain dataset.

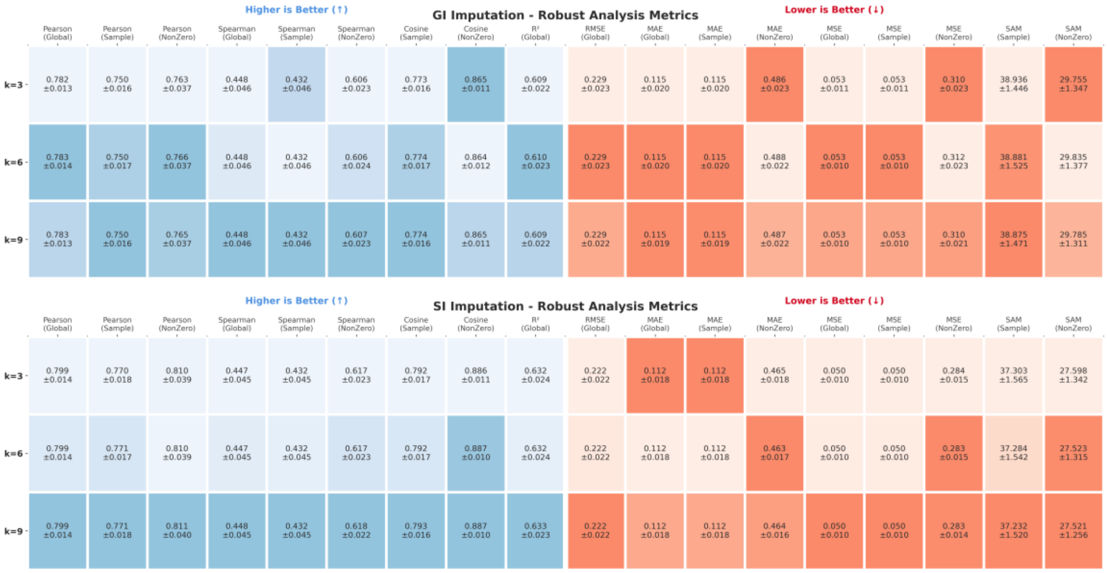


**Supplementary Figure 5.** Comparison of CPS performance for gene and spatial imputation on teacher graphs with neighborhood sizes (k) of 3, 6, and 9. CPS demonstrates robust performance across all evaluated neighborhood sizes.

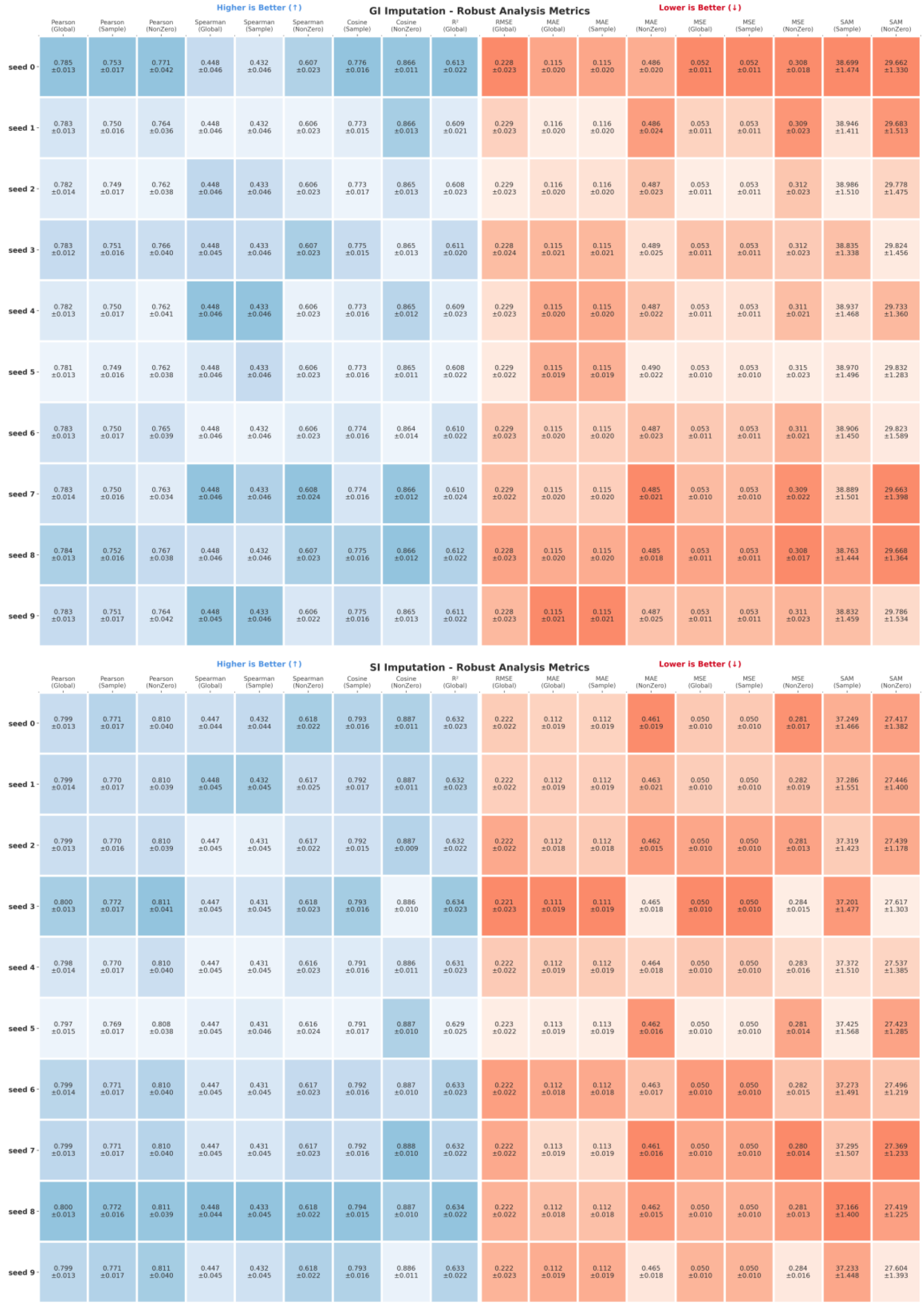


**Supplementary Figure 6.** Comparison of CPS performance in gene and spatial imputation tasks with random seeds from 0 to 10. CPS demonstrates consistent performance across all random seeds.

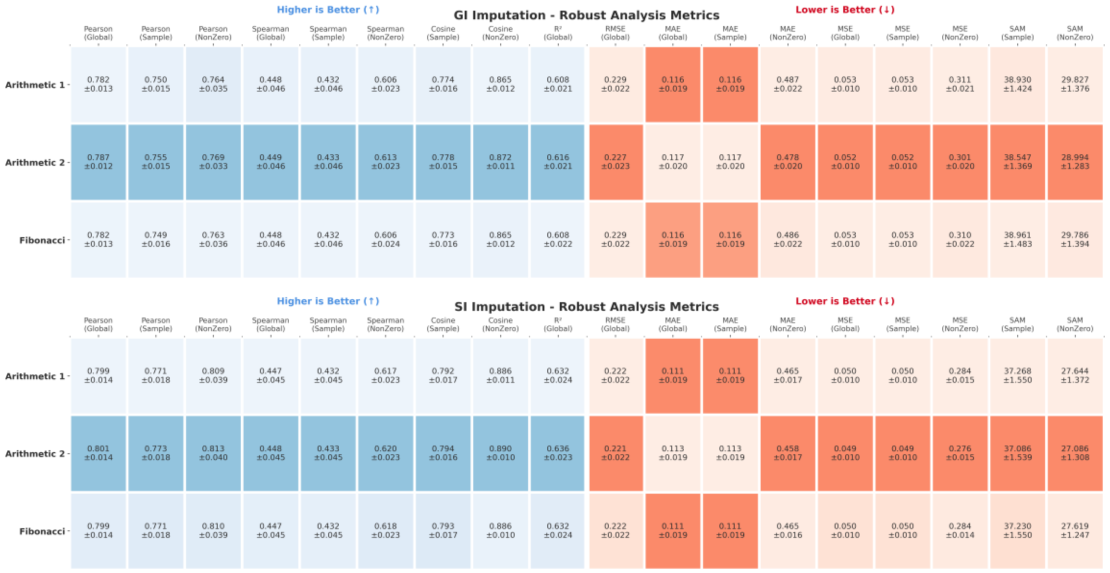


**Supplementary Figure 7.** Comparison of performance metrics in gene and spatial imputation tasks for multi-scale niche attention constructed with arithmetic sequences (1 and 2) and Fibonacci sequences. The models built with arithmetic sequences achieve better performance.

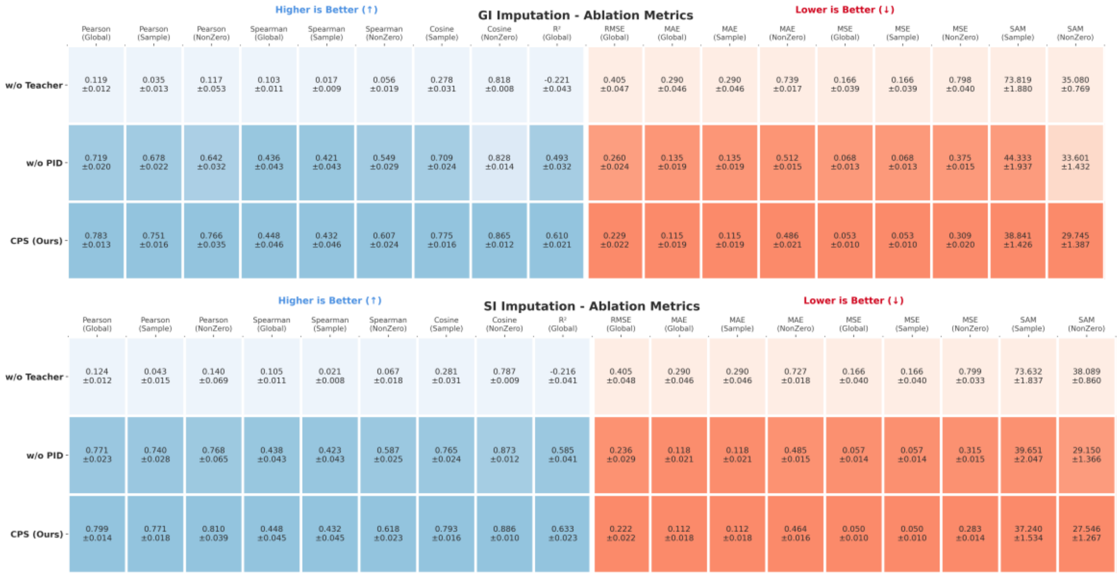


**Supplementary Figure 8.** Comparison of gene imputation and spatial imputation metrics for CPS, ablated teacher network, and ablated PID loss. CPS achieves the best performance, and ablating the teacher network has a greater impact on the metrics than ablating the PID loss.

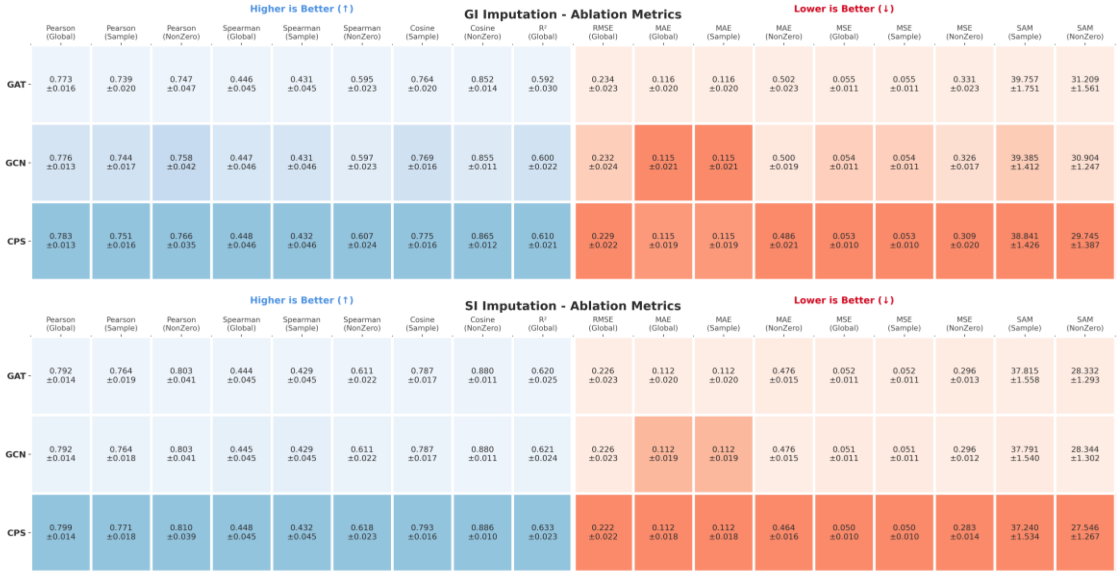


**Supplementary Figure 9.** Comparison of gene imputation and spatial imputation metrics for CPS, and its variants with GCN and GAT replacing the multi-scale niche attention module. CPS achieves the best performance.

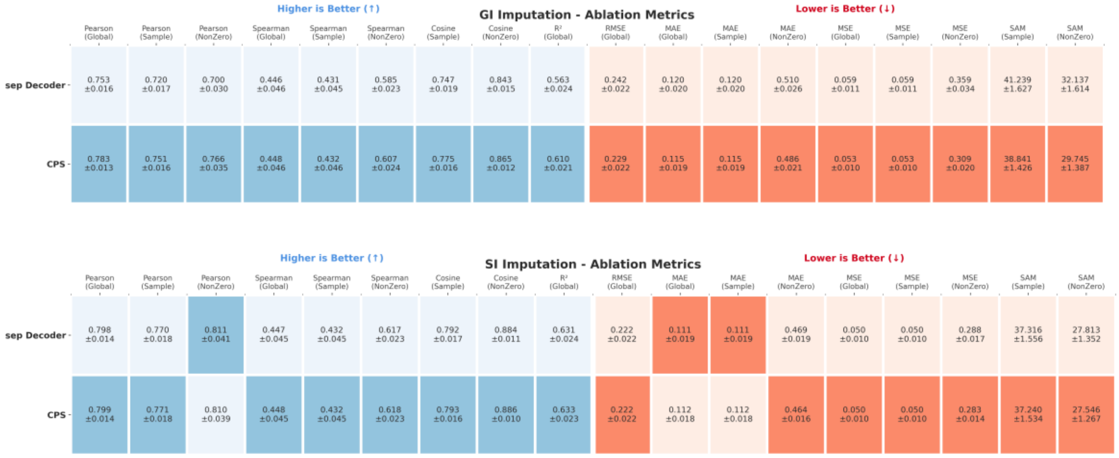


**Supplementary Figure 10.** Comparison of spatial imputation and gene imputation metrics for CPS with a separate decoder and a frozen decoder. The model with a frozen decoder achieves better performance.

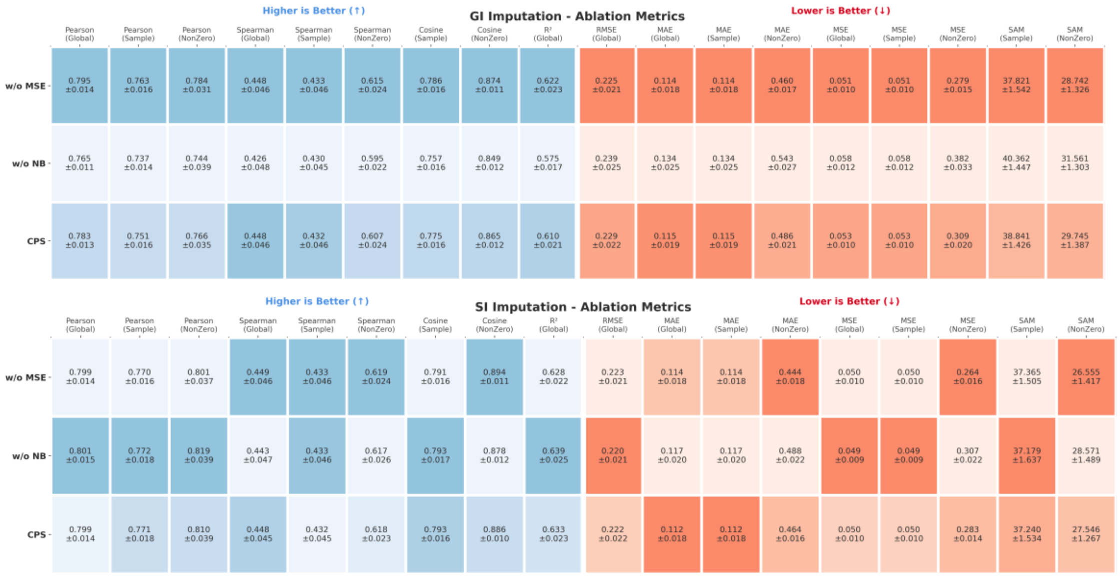


**Supplementary Figure 11.** Comparison of gene and spatial imputation metrics for CPS, CPS without MSE loss, and CPS without NB loss. In gene imputation, the model without MSE loss achieves slightly better performance. In spatial imputation, the model without NB loss yields better results. The standard CPS, utilizing both MSE and NB losses, achieves balanced performance across both tasks.

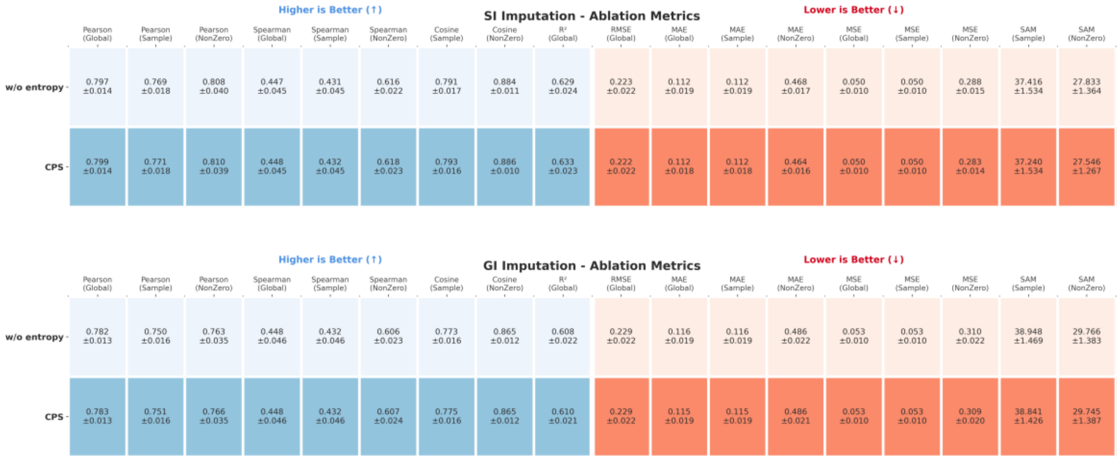


**Supplementary Figure 12.** Comparison of gene and spatial imputation performance for CPS and CPS without entropy regularization loss. The standard CPS achieves better performance in both tasks.

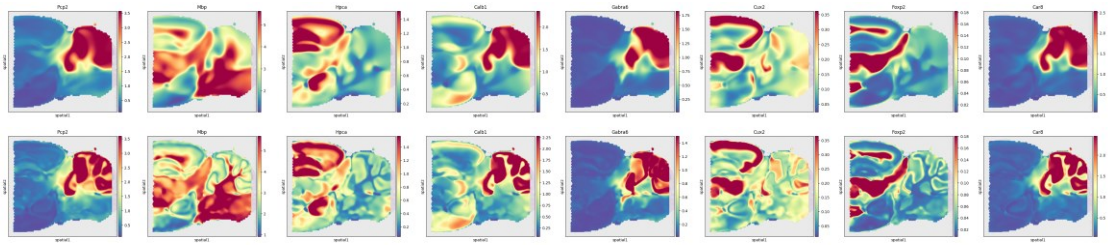

**Supplementary Figure 13.** Comparison of CPS using implicit neural representation (INR, bottom) and multilayer perceptron (MLP, top) on several marker genes in the super-resolution task. INR yields sharper and clearer structural reconstructions compared to MLP.

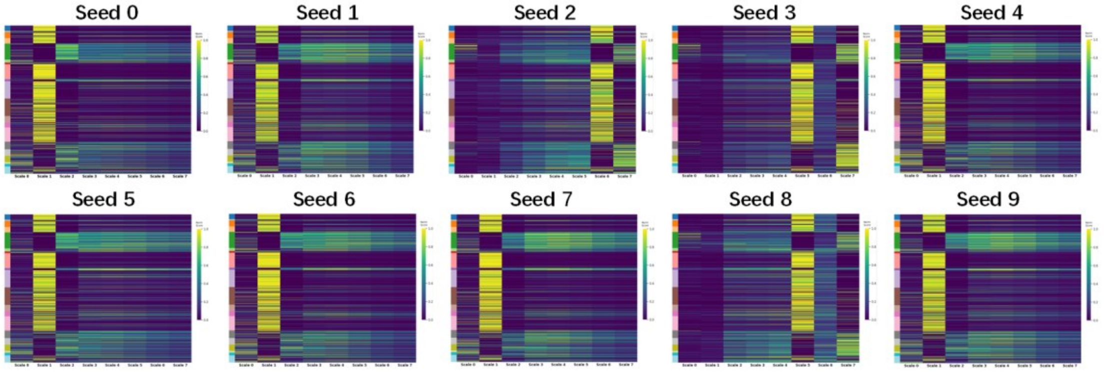


**Supplementary Figure 14.** Sensitivity analysis of the multi-scale niche attention in the CPS teacher network to different random seeds (0-9). The model stably identifies distinct patterns within the tumor, at the tumor edge, and healthy tissue. In 70% of the seeds, long-range attention patterns are observed at the tumor edge and healthy tissue, while short-range patterns dominate within the tumor. The remaining 30% of seeds (2, 3, 8) exhibit long-range attention patterns within the tumor. Overall, our CPS model demonstrates relative stability.

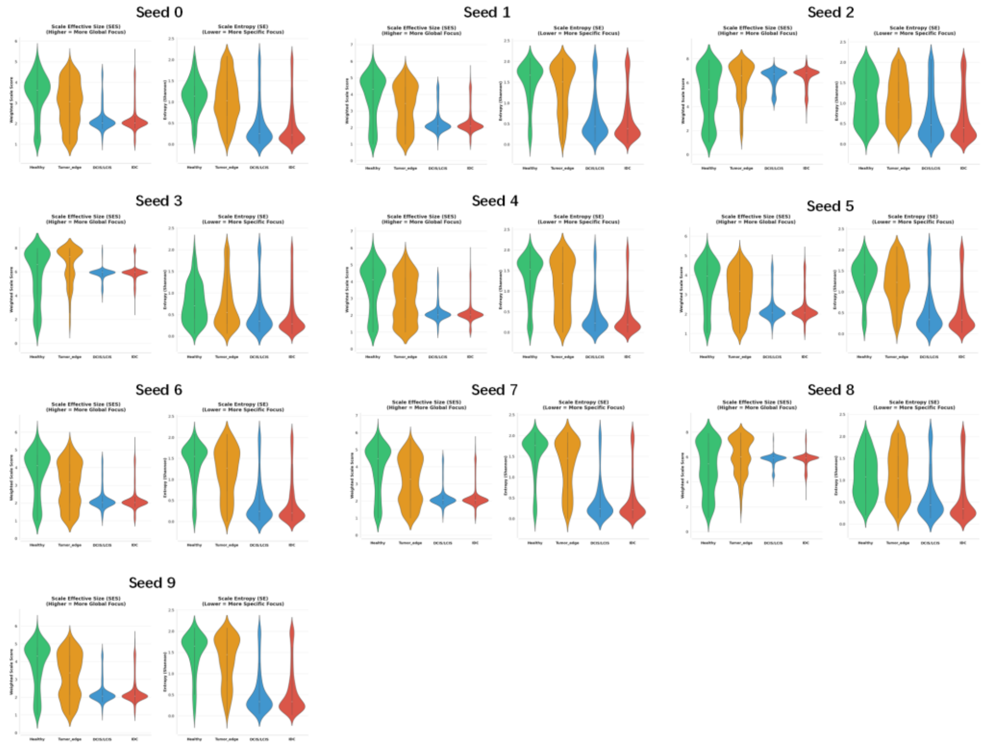


**Supplementary Figure 15.** Sensitivity analysis of the multi-scale niche attention in the CPS teacher network to different random seeds (0-9), visualized by the scale effective size (SES) and scale entropy (SE) computed for each seed. The SES and SE exhibit statistical patterns consistent with the observed attention patterns.

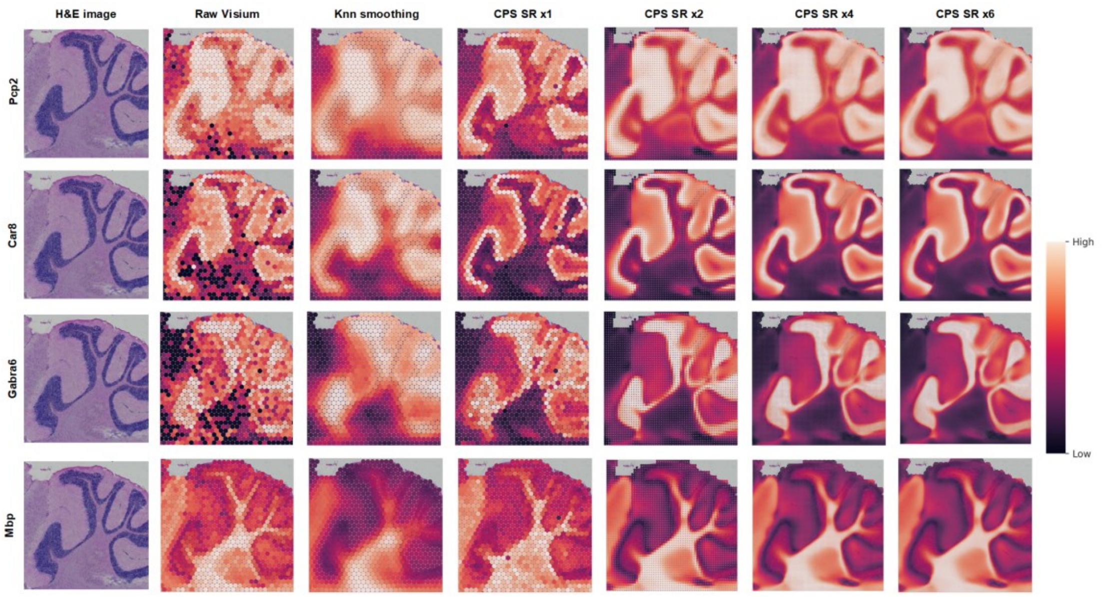


**Supplementary Figure 16.** Visualization of the super-resolution results on marker genes within regions of interest (ROIs). The super-resolved gene expression exhibits sharper boundaries and higher contrast compared to the k-NN smoothed results.

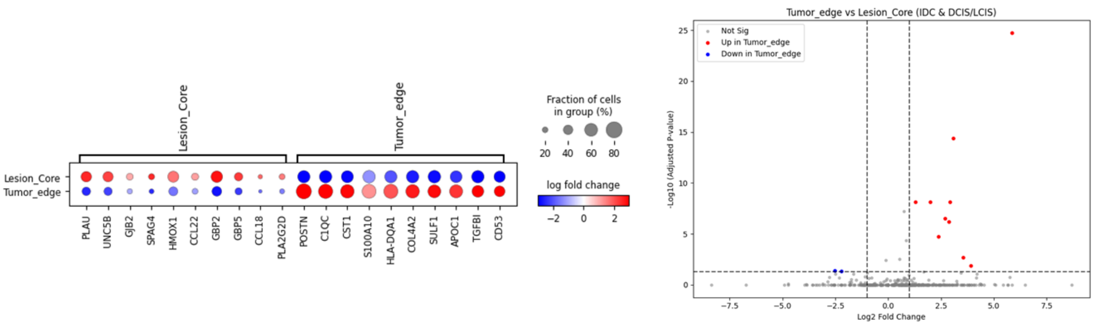


**Supplementary Figure 17.** Differential expression analysis on the HBC dataset using raw Visium data for Tumor edge versus Lesion Core (IDC and DCIS/LCIS) regions. The analysis reveals limited differential expression, resulting in the identification of a small number of significant genes.

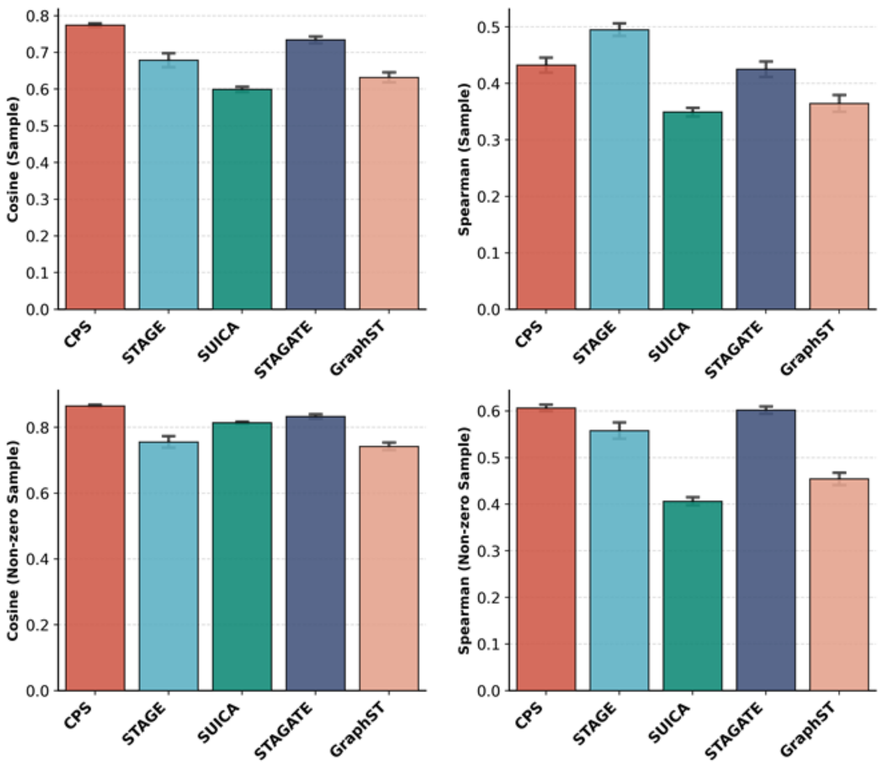


**Supplementary Figure 18.** Comparison of CPS spatial imputation with baseline methods using cosine similarity, and Spearman correlation (sample-level and non-zero value comparisons). CPS achieves superior performance on all metrics except for the sample-level Spearman correlation.

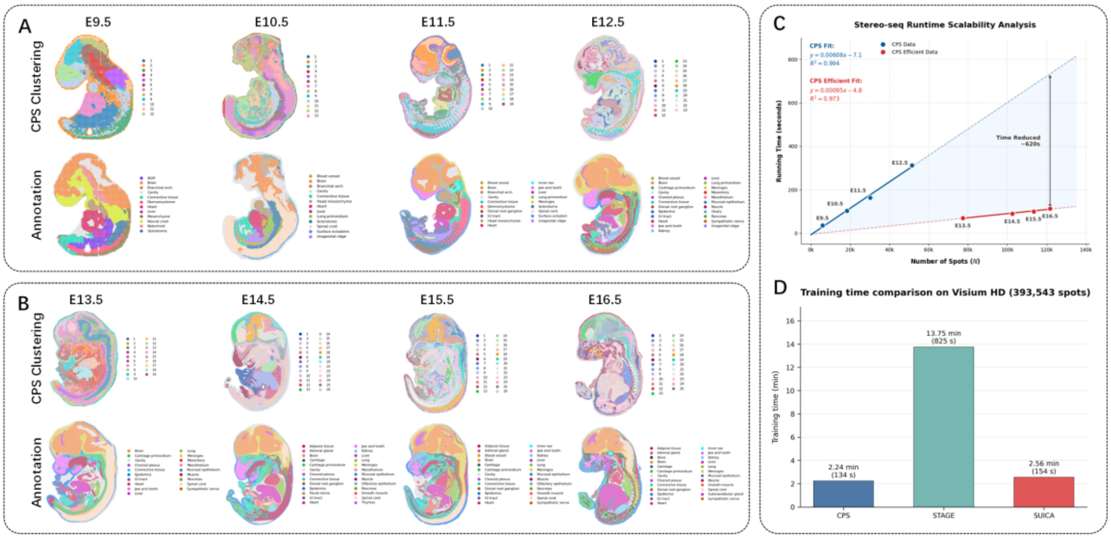


**Supplementary Figure 19.** Scalability and efficiency analysis of CPS on large-scale datasets. A. Clustering performance on Stereo-seq mouse embryo datasets (E9.5 to E12.5) using standard CPS graph tokens, showing high agreement with anatomical annotations. B. Clustering performance on late-stage embryos (E13.5 to E16.5) using the Efficient Mode (pre-processed graph tokens), demonstrating robustness to massive data scales. C. Runtime scalability analysis. The Efficient Mode (Red) significantly reduces computational time compared to the standard mode (Blue), achieving a 6 times speedup with linear complexity (N=20k to 140k spots). D. Training time comparison on a Visium HD dataset (N=393,543). CPS achieves the lowest training time (2.24 min) compared to SUICA and STAGE.

#### Supplementary Tables

**Supplementary Table 1.** Hyperparameters setting of CPS model

| Hyper-Parameters | | Default Setting |
|---|---|---|
| **Teacher Network** | k neighbors | 6 for Visium, 8 for grid base Visium HD and Stereo-seq |
| | neighbor radius | 150/300 for Visium (only for RKNN, in KNN method can be set to 0) |
| | k_list | from 0 to 7, 0 is spot-self |
| | attention head | 4 |
| | latent dim | 64 |
| | dropout | 0.2 |
| **Student Network** | sigma | 1 (higher may introduce noise) |
| | Fourier frequency | 32 (at least 8, higher is redundance) |
| | INR layers | 3 |
| | INR band | 256 |
| | latent dim | 64 |
| | dropout | 0.2 |
| | Coords dim | 2 |
| **Decoder** | latent dim | 64 |
| | middle dim | [256, 512, 1024] |
| | dropout | 0.2 |
| **Loss weights** | teacher Rec | 1.0 |
| | student Rec | 1.0 |
| | distilled weights | 1.0 |
| | entropy freedom | 50% |
| | entropy regularization | 0.05 (higher is fine) |
| **Training** | learning rate | 1e-3 |
| | weight decay | 1e-4 |
| | batch size | 256 (only used in efficient fit) |
| | training epochs | 1e3 (efficient fit loss backwards > 1e3 times) |

For more details, please refer to our reproducibility notebook on: https://github.com/tju-zl/CPS.
